# Characterization of Sex-Based Differences in Integrin-Mediated Endothelial Cell Adhesion to Bioactive Hydrogels

**DOI:** 10.1101/2025.06.02.656802

**Authors:** Abbey Nkansah, Aashee Budhwani, Nicholas Grammer, Nicolai Ang, Ashauntee Fairley, Sabrina Pietrosemoli Salazar, Aarushi Chowdary Yedalla, Adhwita Anand, Stephanie Seidlits, Josephine Allen, Elizabeth Cosgriff-Hernandez

**Author notes:** Address for correspondence: Elizabeth Cosgriff-Hernandez, Ph.D., Department of Biomedical Engineering, The University of Texas at Austin, 107 W. Dean Keeton, BME Building, Room 3.503D, Austin, TX 78712, USA.

## Abstract

Endothelialization promotes thromboresistance in blood-contacting devices, but biomaterial designs often overlook sex differences in endothelialization processes. In this study, we elucidated sex differences in endothelial cell-material interactions through investigation of the integrin-ligand interplay with biomaterial substrates and the corollary effects on cell adhesion and spreading. First, integrin expression of human coronary artery endothelial cells (HCAECs) was characterized for age-matched donors (3 male, 3 female). Sex-based differences in integrin expression were identified, with notably higher α2β1, α5β1, and αVβ3 expression in female cells and higher α1β1 expression in male cells. On polyethylene glycol (PEG)-based hydrogel incorporating collagen or gelatin, female cells showed increased attachment on stiff substrates as compared to male HCAECs, likely driven by increased α2β1, α5β1, and αVβ3 expression in female cells. Collectively, these results demonstrate sex-biased endothelial cell responses to bioactive hydrogels mediated by integrin interactions and highlight the importance of incorporating biological sex as a design variable in the development of blood-contacting biomaterials.

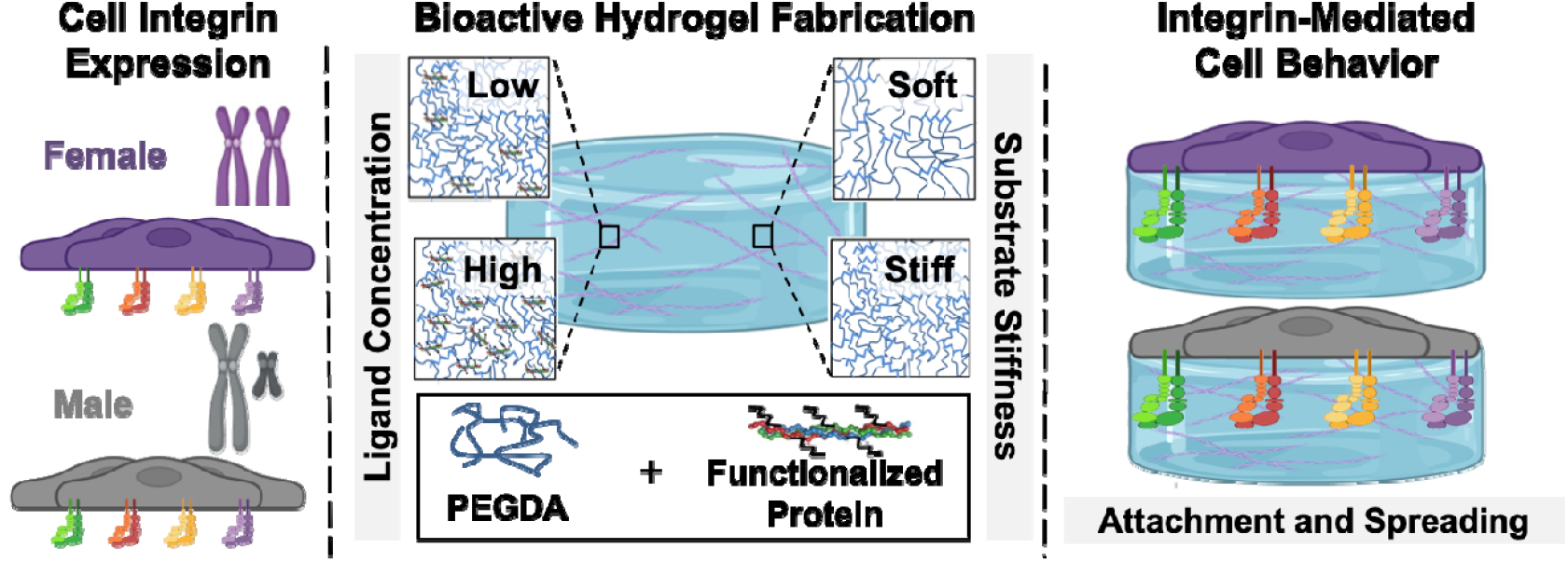

## 1. Introduction

Endothelialization has been widely adopted as the best strategy to prevent thrombotic failure in synthetic blood-contacting devices as endothelial cells play a critical role in long-term hemostatic regulation.^1-3^ Endothelial cell interactions that drive graft coverage are in part dictated by integrin expression by the endothelial cells, substrate ligand identity and ligand concentration, and substrate stiffness as these mechanisms can influence adhesion strength and conferral of thromboresistant phenotypes. Despite concerted research efforts over several decades, scientists simply have not replicated the necessary cellular cues to promote full graft endothelialization. In addition, intrinsic sex-based differences in endothelial cell behavior may further confound these efforts due to variability in thromboresistant phenotypes and contribute to disparities in blood-contacting device success.^4-6^ However, there is little known about the role of sex in integrin-mediated endothelialization processes that underpin most efforts to design thromboresistant devices.

Endothelial cell interactions with extracellular matrix ligands govern cell adhesion, spreading, and migration for sustained thromboresistance.^3, 7^ Extracellular matrix proteins (e.g., collagen, fibronectin, laminin) and proteins/peptides derived from the extracellular matrix (e.g., gelatin, RGD) have been extensively explored as coatings for blood-contacting devices to impart bioactivity and promote endothelialization.^3, 8-10^ However, limited studies have been performed to characterize integrin-mediated cell behavior and hemostatic regulation based on sex.^4^ To this end, we have developed a bioactive polyethylene glycol (PEG)-based hydrogel platform with facile incorporation of bioactive cues that can be used to decouple the effects of ligand concentration, ligand identity, and substrate stiffness to better isolate sex-differences in cell material interactions.^10^ Incorporation of ligands such as collagen and gelatin conjugated to PEG-hydrogels provides an array of critical integrin targeting motifs that constitute the ECM. Post et al. determined that differences in human umbilical vein endothelial cell (HUVEC) integrin expression on bioactive hydrogels incorporating collagen and/or gelatin leads to differences in attachment and hemostatic regulation.^3^ These integrin-targeting hydrogels provide new tools to unravel critical sex-based differences in integrin-mediated endothelialization processes.

In this study, we elucidated sex differences in endothelial cell-material interactions through investigation of the integrin-ligand interplay with biomaterial substrates and the corollary effects on cell adhesion and spreading. To understand the underlying mechanisms governing sex-biased endothelial cell behavior, we first assessed integrin expression differences (α1, α2, α5, αv) between age-matched male and female human coronary artery endothelial cells (HCAECs). These were selected based on the established role of α1β1 and α2β1 mediated attachment to collagen and α5β1 and αvβ3 mediated attachment to fibronectin, gelatin, and RGD-containing substrates.^3, 8, 11-15^ The sex-based integrin expression differences were then utilized to elucidate cell behavior on integrin-targeting hydrogel substrates. A tunable, bioactive PEG-based hydrogel platform was used to systematically investigate the effect of ligand concentration, ligand identity, and substrate stiffness. As limited studies have been conducted to elucidate sex dimorphisms in endothelial cell behavior, this investigation will serve as a guide for researchers to elucidate how sex-based differences in integrin engagement impacts cell-material interactions. This work will also provide a framework for future investigations of other patient variables beyond sex differences (e.g. age, ancestry, hormone) to ensure equal health outcomes regardless of patient demographics.

## 2. Results and Discussion

Integrin expression differences between male and female cells were first characterized using flow cytometry. Regardless of sex, it was observed that *α*1 expression was lowest, followed by *α*V, *α*2, *α*5 expression, respectively (**Figure 1**). Significant differences in integrin expression between male and female cells were observed for each integrin subunit. Male cells displayed a significant increase in expression of *α*1; whereas, female cells displayed an increased expression of α2, α5, and αv. The apparent increased female cell expression of αv subunit is consistent with previous studies elucidating sex-based difference in αvβ3 integrin expression in male and female cells originating from skeletal muscle microvasculature.^5^ These results indicate a sex bias in HCAEC integrin expression of *α*1β1 and α2β1 which are integral to attachment to collagen and α5β1 and αvβ3 which are involved in attachment to gelatin and fibronectin.

**Figure 1.**
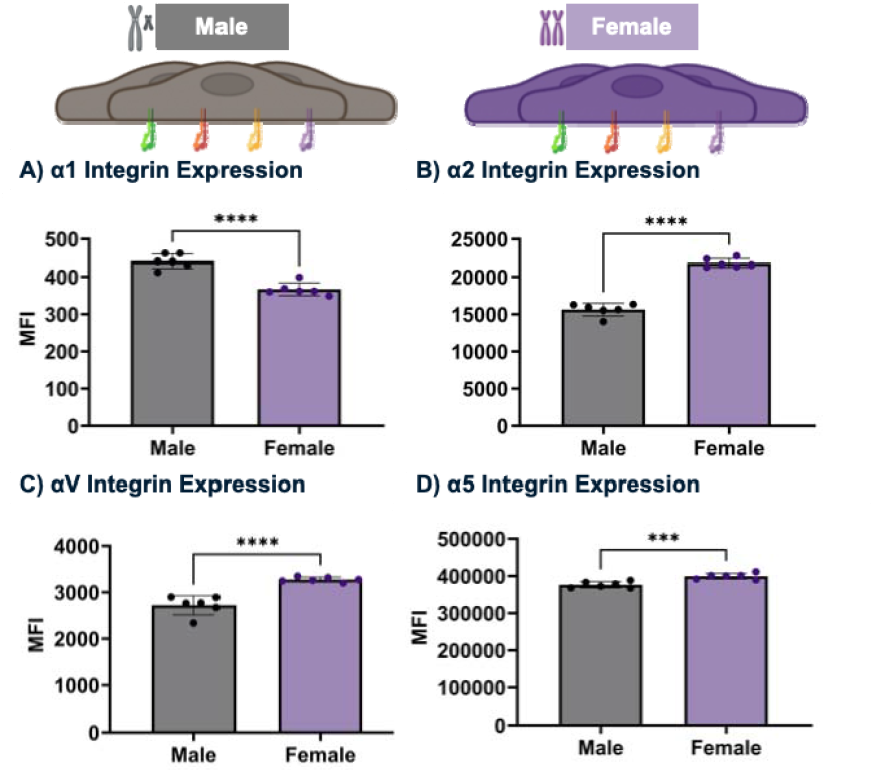
Sex differences in relative abundance of key integrin subunits involved in endothelial cell attachment to extracellular matrix proteins. Relative fluorescence was measured using flow cytometry using antibodies A) anti-α1, B) anti-α2, C) anti-αV, and D) anti-α5. Two biological replicates were performed for each donor (n = 3 per sex). Data for all comparisons represents the average of each biological replicate and standard deviation (*** = p < 0.001; **** = p < 0.0001)

Adhesion and spreading measurements of HCAECs cultured on bioactive hydrogels were used as an initial first step in evaluating sex-based differences in integrin-mediated cell-material interactions. We employed a bioactive PEG-based hydrogel platform that incorporated collagen or gelatin at different ligand concentrations and different substrate stiffnesses. Collagen was chosen for its prominence in the basement membrane with integrin binding motifs critical to endothelial cell adhesion and phenotypes via interaction with α1β1 and α2β1 integrins.^16-18^ It is also commonly utilized in hydrogels and other ECM-based platforms in attempts to recapitulate innate cellular environments.^19-21^ Gelatin, the denatured form of collagen, was selected to elucidate interactions with α5β1 and αvβ3 integrins which are targeted in biomaterial design using RGD and CRRETAWAC peptides or ECM proteins such as fibronectin or laminin.^8, 22-26^ These integrin binding motifs are critical in modulating endothelial cell function and hemostatic regulation on blood-contacting devices.^8^ Previous literature was used to select a ligand concentration range that promotes integrin engagement without exceeding ligand signal saturation that could dampen cell spreading effects, enabling detection of sex-specific differences in integrin engagement.^27^ Two ligand concentrations (Low: 0.1 and High: 2.0 mg/mL) were used to investigate the interplay of ligand availability and integrin expression in regulating cellular responses, encompassing conditions both below and at ligand saturation levels.^27^ The stiffnesses utilized in this study (Soft: 10 kPa and Stiff: 110 kPa) were selected to elucidate significant differences in cell spreading. ECs have been shown to exhibit enhanced spreading on stiffer substrates, given their increased ability to apply traction forces on the underlying matrix. Early investigations of matrix stiffness with hydrogels have been largely focused on soft matrices (<10 kPa) for independent tuning of protein concentration and substrate stiffness to promote angiogenic cues.^28-29^ Stiffer substrates were selected to span closer to physiologically relevant ranges for vascular tissues.^30^ A factorial design was then utilized to investigate the individual and synergistic effects of ligand identity, ligand concentration, and substrate stiffness.

First, sex differences in HCAEC adhesion and spreading on PEG-Collagen and PEG-Gelatin hydrogels were evaluated after 24 hours at low and high protein concentrations (0.1 mg/ml and 2 mg/ml). Across both bioactive hydrogels, increases in cell attachment and spreading with increased protein concentrations were observed regardless of sex. These trends are consistent with previous studies demonstrating increased ligand density leads to increased attachment and spreading up to a point of ligand saturation.^27^ Female cells on PEG-collagen hydrogels demonstrated increased attachment on substrates with low ligand concentrations in comparison to male cells cultured under similar conditions but no differences was observed at higher ligand concentration (**Figure 2A, C)**. As α2β1 expression was greater for female cells over male cells, with four orders of magnitude increase over α1β1 expression, this likely indicates that of the investigated integrins, α2β1 is the dominant integrin mediated cell adhesion to collagen and that increased expression in females leads to greater differences at low protein concentration. At higher protein concentration, the interplay with other adhesion motifs such as syndecan-1 may potentially offset this sex-based difference in integrin-mediated attachment and spreading.^31^ No significant sex-based differences in endothelial cell spreading were observed with either ligand concentation. On PEG-gelatin hydrogels, no significant differences in attachment and spreading were observed at lower concentrations. At higher concentrations of gelatin, significant increases in female cell attachment and spreading were observed (**Figure 2B, D**). It was hypothesized that this could be due to increased α5β1 and αVβ3 expression for female cells over male cells, leading to increased opportunities for binding affinity for female cells.

**Figure 2.**
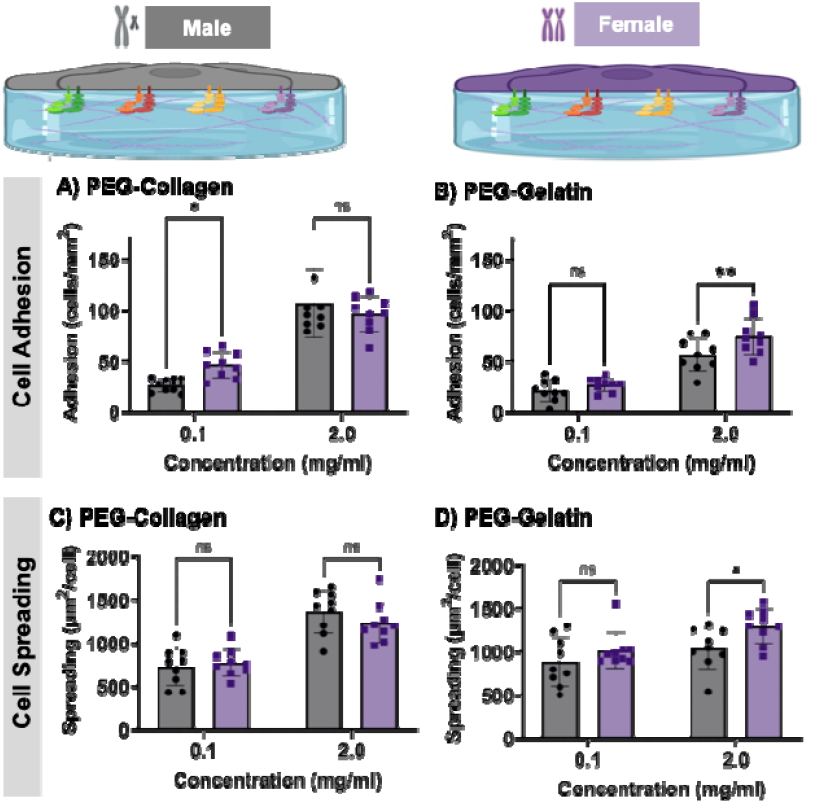
Effects of ligand identity and concentration on cell attachment and spreading of female and male endothelial cells cultured on bioactive hydrogels. Attachment and spreading on A,C) PEG-collagen and (B,D) PEG-gelatin (n = 3 specimen per donor). Mean and standard deviation for each specimen is represented (ns = not significant; * = p < 0.05, ** = p < 0.01)

In comparing the effect of ligand identity, it was noted that male cells displayed increased attachment and spreadin gon PEG-collagen substrates as compared to PEG-gelatin; however, there was no significant difference observed with female cells (**Figure S2**). It was hypothesized that this could be due to increased α5β1 and αVβ3 expression for female cells over male cells, leading to increased opportunities for binding to gelatin ligands. It was observed that low ligand concentrations demonstrated greater sex-based effects in attachment to PEG-collagen substrates in comparison to PEG-gelatin substrates (**Figure 2A, B**). Whereas, greater sex-based effects were noted on PEG-gelatin than PEG-collagen substrates at high concentrations with increased attachment and spreading for female cells compared to male cells.

To elucidate sex differences in cell behavior based on substrate stiffness, collagen and gelatin were incorporated into PEG hydrogels (concentration = 2.0 mg/mL) at different PEG hydrogel crosslink densities. Low PEGDA macromer molecular weight (3.4 kDa) were utilized to fabricate stiff substrates (∼110 kDa) and high PEGDA macromer molecular weight (20 kDa) were used to fabricate softer substrates (∼10 kPa), providing a 90% decrease in modulus while maintaing constant protein concentrations (**Figure S1**). Overall, stiffness led to an increase in attachment and spreading on both bioactive hydrogel substrates. Neither soft nor stiff PEG-collagen hydrogels demonstrated significant differences in attachment or spreading based on sex (**Figure 3A, C)**. Similarly, there were no distinguishable sex differences in cell attachment or spreading on soft PEG-gelatin hydrogels. We hypothesize that low-modulus substrates may not induce sufficient mechanical cues necessary to induce significant differences integrin-mediated signaling needed to reveal the sex-based integrin expression differences previously described.^27^ However, with increased stiffness, a marked increase in adhesion and spreading for female cells in comparison to male cells was observed on PEG-gelatin hydrogels (**Figure 3B, D)**. It was hypothesized that this was due to increased expression in female cells of α5β1 and αVβ3 motifs. Future studies will characterize sex-based integrin expression differences from HCAECs cultured on a broader range of substrate stiffnesses and ligand concentration to help elucidate the lack of sex-dependent differences observed at low substrate stiffness.

**Figure 3.**
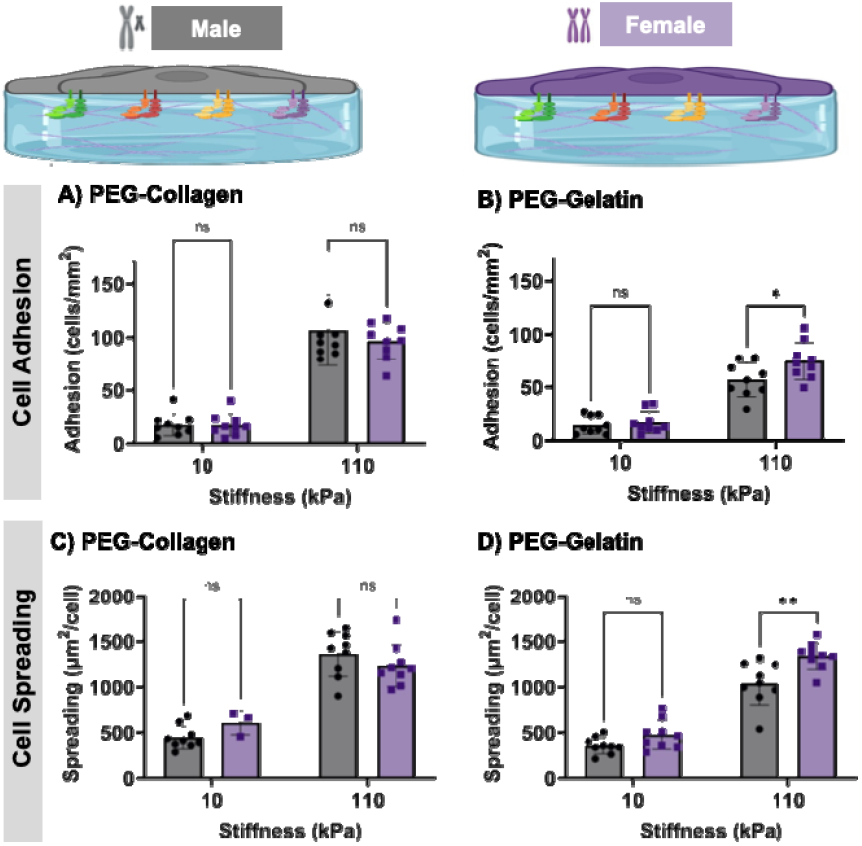
Effects of stiffness on cell attachment and spreading of female and male endothelial cells to bioactive hydrogels (ligand concentration = 2.0 mg/mL). Attachment and spreading on A,C) PEG-collagen (n = 3 specimen per donor) and B,D) PEG-gelatin (n = 3 specimen per donor). Mean and standard deviation for each specimen is (ns = not significant; * = p < 0.05, ** = p < 0.01).

Endothelialization is vital to ensure long-term thromboresistance of blood-contacting devices.^1, 8^ Given the growing evidence of differential cellular responses due to patient variables (e.g., sex, age, ancestry), it is critical to consider these variables in biomaterial designs to ensure equitable patient outcomes. This study provides valuable insights into sex-based differences in endothelial cell integrin expression and behavior on bioactive hydrogels; however, we wanted to note limitations of this first effort to characterize sex-based effects on integrin-mediated endothelialization. First, integrin expression was measured at a single time point following expansion on tissue culture polystyrene which limits our understanding of how expression patterns may evolve over time and influence dynamic cell responses. It is well established that integrin expression is dynamic and responsive to substrate cues.^23, 32-33^ Future studies will focus on characterizing integrin expression in native tissue with fresh harvest of endothelial cells and integrin expression profile changes after culture on each biomaterial substrate. Second, only a narrow range of ligand concentrations and substrate stiffnesses were evaluated, which may not fully capture the range of biomechanical and biochemical conditions present in vivo. Finally, there are a large number of additional patient variables (e.g., sex hormones, age, ancestry, disease state) that can affect integrin expression and endothelial cell behavior, which could be a significant factor in sex-specific responses. The study design was used to minimize the effect of these confounding variables by age-matching donors and limiting hormone exposure (phenol red free media and charcoal-stripped FBS) but is not able to fully eliminate these variables. Overall, this study serves as a first step toward characterizing sex-based differences in integrin expression and serves as a framework for future investigation of the individual and synergistic effects of these variables on integrin-mediated endothelialization processes.

## 3. Conclusions

In this study, we examined sex-based differences in the expression of integrins α1β1, α2β1, α5β1, and αVβ3 —key mediators of cell adhesion through interactions with extracellular matrix proteins like collagen, fibronectin, and gelatin. A tunable, bioactive PEG-based hydrogel platform was then used to systematically investigate sex-based differences in endothelial cell behavior in response to ligand concentration, ligand identity, and substrate stiffness. Integrin expression was determined to be influenced by sex, with female cells displaying increased expression of α2β1, αvβ3, and α5β1 while male cells had increased expression of α1β1. Our findings also demonstrated that endothelial cell adhesion and spreading are influenced not only by the biochemical and biomechanical cues, but also by the biological sex of the donor cells. In particular, sex bias in attachment and spreading were observed on PEG-collagen at low concentrations and on stiffer substrates with higher concentrations for female cells having enhanced cell responses – likely due to increased expression of α2β1 for female cells compared to male cells. Overall, this data underscores the importance of incorporating sex as a biological variable in the design and evaluation of biomaterials for blood-contacting applications. Future work will include utilizing this new knowledge to design integrin-targeting hydrogels to address sex-biased cell behaviors and optimize devices to promote thromboresistant phenotypes.

## Materials and Methods Materials

Reagents were purchased from Sigma-Aldrich and used without further purification unless otherwise noted.

### Cell culture

Three male and three female donor HCAECs were purchased from Promocell. Donors were age-matched, ranging from 50 to 65 years old, to minimize the confounding effect of age on cell behavior. Each donor set was expanded using Endothelial Cell Growth Media (Promocell) and 20% fetal bovine serum (FBS). FBS was charcoal stripped to remove lipophilic molecules, including hormones, that can interfere with experimental studies. Cells were expanded in media with phenol red because of the increased proliferation rate; however, the media was replaced with phenol red-free media 24 hours before each study to limit estrogenic effects from the phenol red. All cultures were grown in tissue culture polystyrene flasks incubated at 37^°^C with 5% CO_2_. Media was exchanged every 2-3 days.

### Flow Cytometry

Differences in integrin expression between the male and female HCAECs was determined using antibody staining for flow cytometry. HCAECs were cultured in tissue culture polystyrene flasks to 90% confluency. The cells were then lifted from the flask using Accutase and spun down at 220 rcf followed by resuspension in flow buffer (2% FBS in phosphate buffered saline (PBS)) at a concentration of 10^5^ cells/ml. Each sample was stained with fixable viability dye 450 (BD Biosciences) on ice for 20 minutes to selectively analyze viable cells. After washing samples with 3 mL of flow buffer, cells were centrifuged and resuspended in 500 μL of flow buffer. Samples were titrated with their respective integrin antibodies (anti-α1, anti-α2, anti-α5, and anti-αV) in 0.5 ug increments on ice to identify signal saturation. Each sample was then washed with 3 mL of flow buffer and spun down. Cells were resuspended in 400 uL 4% paraformaldehyde for fixation for 15 minutes. Blank samples of cells without antibodies underwent the similar protocols. Samples were then rinsed with 3 mL flow buffer, spun down, and resuspended in 1 mL flow buffer. An Attune Nxt Acoustic Focusing flow cytometer was used to evaluate surface expression of integrins. Relative fluorescence of each integrin was assessed with each donor tested on two separate occasions (biological replicate). For each integrin tested, samples were run in triplicate (n = 3 per donor). Fluorescence intensity was analyzed using FlowJo software. The average of each biological replicate

### Synthesis of Polyethylene Glycol Diacrylate

Polyethylene glycol diacrylate (PEGDA) was synthesized using the protocol from Browning et al. with minor alterations.^34^ Briefly, a solution of PEG (molecular weight 3.4 kDa or 20 kDa; 1 equivalent) was prepared in anhydrous dichloromethane under nitrogen. Triethylamine (2 molar equivalents) was added dropwise to the PEG solution, followed by dropwise addition of acryloyl chloride (4 molar equivalents). The solution was allowed to mix for 24 hours. The reaction solution was then neutralized with potassium bicarbonate (8 mole equivalents) and dried with sodium sulfate. The polymer was precipitated in cold ether, vacuum filtered, and then dried under vacuum. Synthesis of PEGDA was confirmed using proton nuclear magnetic resonance (^1^H-NMR) spectroscopy. Proton NMR spectra were recorded on a Mercury 300 MHz spectrometer using a TMS/solvent signal as an internal reference. ^1^H-NMR (CDCl_3_): 3.6 ppm (m, -OC*H*_*2*_C*H*_*2*_*-*), 4.3 ppm (t, -C*H*_*2*_OCO-) 6.1 ppm (dd, -C*H*=CH_2_), 5.8 and 6.4 ppm (dd, - CH=C*H*_*2*_). Polymers with percentage conversions of hydroxyl to acrylate end groups over 85% were used in this work.

### Bioactive Hydrogel Fabrication and Characterization

Bioactivity was imparted to PEGDA (3.4 or 20 kDa) hydrogels through incorporation of either collagen (rat tail type I) or gelatin functionalized with an acrylamide-isocyanate linker (functionalization of ∼10% of available lysines).^34^ PEG-collagen hydrogels were prepared by dissolving PEGDA and functionalized collagen (0.1-2.0 mg/mL) in 20 mM acetic acid. PEG-gelatin (type B) hydrogels were fabricated by dissolving 10 wt% PEGDA and functionalized gelatin (0.1-2.0 mg/mL) in deionized water. A photoinitiator solution of Irgacure 2959 (10 wt/vol% solution in 70 vol% ethanol) was added to precursor solutions at a final concentration of 0.1 wt/vol%. Hydrogel sheets were fabricated by pipetting precursor solutions between 1.5 mm spaced glass plates and curing on a UV transilluminator (UVP, 25 watt, 365 nm) for 6 minutes on both sides. Blank hydrogel specimens devoid of protein were prepared as a negative control with the same protocol.

### Endothelial Cell Adhesion and Spreading

Hydrogel disc specimens (D = 6 mm) were punched from hydrogel sheets with and without protein and placed into a 48 well plate. HCAECs were seeded at 10,000 cells/cm^2^ on hydrogel discs (n = 4 for donor) and allowed to attach for 24 hours. Cells were then washed twice with PBS to remove non-adherent cells. Specimens were then fixed with 3.7% glutaraldehyde and stained with rhodamine phalloidin (actin/cytoskeleton) and SYBR green (DNA/nucleus). Specimens were imaged (3 images per specimen and *n* = 3 specimens per donor for a total of 9 images per donor) using a fluorescence microscope (Nikon Eclipse TS100) to quantify cell adhesion and spreading. The average adhesion and spreading of three specimen images were reported. Cell adhesion was assessed by manually counting the number of SYBRGreen-stained nuclei for each image. The magic wand tool in ImageJ was utilized to quantify average cell spreading of rhodamine phalloidin-stained images. The tolerance of the tool was adjusted to highlight extracellular regions to determine the average pixels per cell. Pixels were then converted to microns to evaluate the average area per cell.

### 2.5. Statistical Analysis

The data for all other measurements are displayed as mean ± standard deviation. Statistical comparisons were made using an analysis of variation (ANOVA) comparison utilizing Tukey’s post-hoc analysis for parametric data. Computations were performed using GraphPad Prism version 10 at the significance levels of p < 0.05.

