## Supplemental Information for "Characterization of Sex-Based Differences in Integrin-Mediated Endothelial Cell Adhesion to Bioactive Hydrogels"

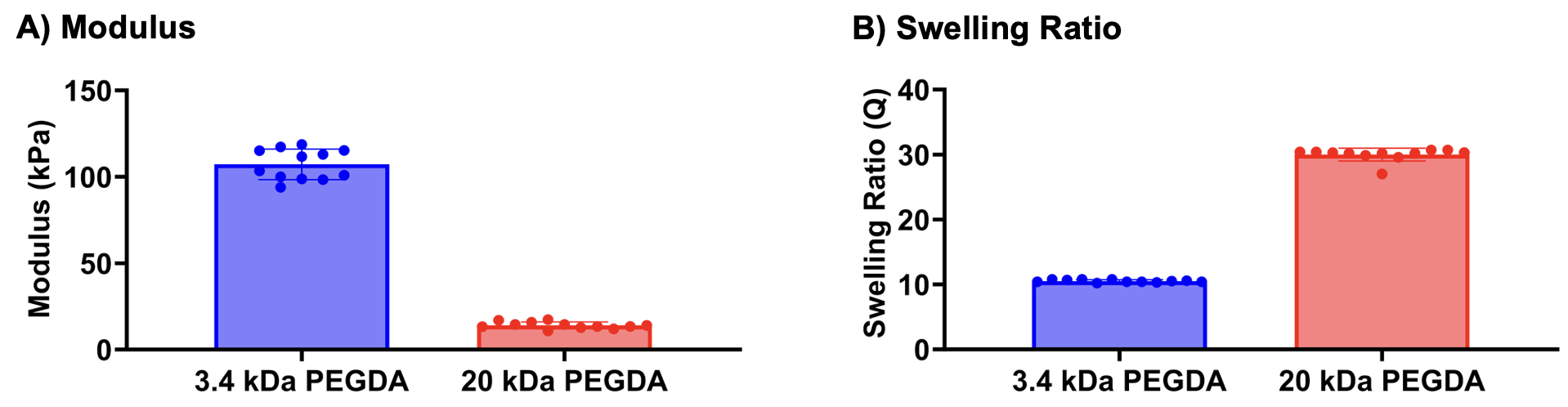

**Figure S1:** Effects of molecular weight on modulus and swelling ratio. n = 12; measurements are expressed as mean average ± standard deviation.

**Table S1:** Calculated p-Values from ANOVA Analysis of the Individual and Combined Effects of protein type, sex, concentration, and stiffness on cell adhesion and spreading to PEG-collagen and PEG-gelatin.

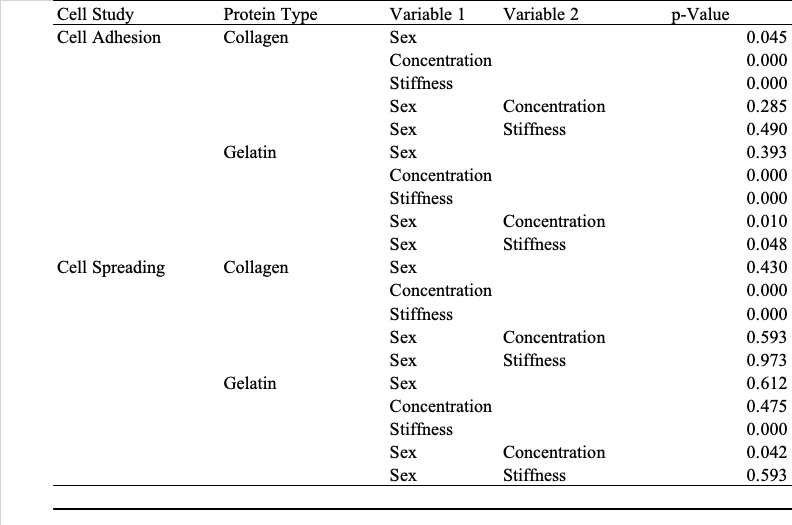

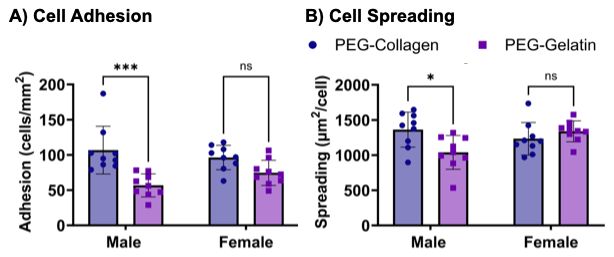

**Figure S2:** Effects of ligand identity on cell attachment and spreading of female and male endothelial cells to bioactive hydrogels (ligand concentration = 2.0 mg/mL). A) Attachment and B) spreading on PEG-collagen and PEG-gelatin (n = 3 specimen per donor). Mean and standard deviation for each specimen is (ns = not significant; * = p < 0.05, *** = p< 0.001).
